# A cross-disorder MR-pheWAS of 5 major psychiatric disorders in UK Biobank

**DOI:** 10.1101/634774

**Authors:** Beate Leppert, Louise AC Millard, Lucy Riglin, George Davey Smith, Anita Thapar, Kate Tilling, Esther Walton, Evie Stergiakouli

## Abstract

Psychiatric disorders are highly heritable and associated with a wide variety of social adversity and physical health problems. Using genetic liability (rather than phenotypic measures of disease) as a proxy for psychiatric disease risk can be a useful alternative for research questions that would traditionally require large cohort studies with long-term follow up.

Here we conducted a hypothesis-free phenome-wide association study in about 300,000 participants from the UK Biobank to examine associations of polygenic risk scores (PRS) for five psychiatric disorders (major depression (MDD), bipolar disorder (BP), schizophrenia (SCZ), attention-deficit/ hyperactivity disorder (ADHD) and autism spectrum disorder (ASD)) with 23,004 outcomes in UK Biobank, using the open-source PHESANT software package.

There was evidence after multiple testing (p<2.55×10^−06^) for associations of PRSs with 226 outcomes, most of them attributed to associations of PRS_MDD_ (n=120) with mental health factors and PRS_ADHD_ (n=77) with socio-demographic factors. Among others, we found strong evidence of associations between a 1 standard deviation increase in PRS_ADHD_ with 1.1 months younger age at first sexual intercourse [95% confidence interval [CI]: −1.26,−0.94]; PRS_ASD_ with 0.01% reduced lower erythrocyte distribution width [95%CI: −0.013,-0.007]; PRS_SCZ_ with 0.98 odds of playing computer games [95%CI:0.976,0.989]; PRS_MDD_ with a 0.11 points higher neuroticism score [95%CI:0.094,0.118] and PRS_BP_ with 1.04 higher odds of having a university degree [95%CI:1.033,1.048].

We were able to show that genetic liabilities for five major psychiatric disorders associate with long-term aspects of adult life, including socio-demographic factors, mental and physical health. This is evident even in individuals from the general population who do not necessarily present with a psychiatric disorder diagnosis.

**AUTHOR SUMMARY:** Psychiatric disorders are associated with a wide range of adverse health, social and economic problems. Our study investigates the association of genetic risk for five common psychiatric disorders with socio-demographics, lifestyle and health of about 330,000 participants in the UK Biobank using a systematic, hypothesis-free approach. We found that genetic risk for attention deficit/hyperactivity disorder (ADHD) and bipolar disorder were most strongly associated with lifestyle factors, such as time of first sexual intercourse and educational attainment. Genetic risks for autism spectrum disorder and schizophrenia were associated with altered blood cell counts and time playing computer games, respectively. Increased genetic risk for depression was associated with other mental health outcomes such as neuroticism and irritability. In general, our results suggest that genetic risk for psychiatric disorders associates with a range of health and lifestyle traits that were measured in adulthood, in individuals from the general population who do not necessarily present with a psychiatric disorder diagnosis. However, it is important to note that these associations aren’t necessary causal but can themselves be influenced by other factors, like socio-economic factors and selection into the cohort. The findings inform future hypotheses to be tested using causally informative designs.

## INTRODUCTION

Family and twin research as well as large-scale genome-wide association studies (GWAS) have shown that psychiatric disorders are highly heritable (1) and that genetic risks for psychiatric disorders also are associated with socio-economic factors, physical health outcomes as well as other psychiatric disorders (2–5). Using genetic liability (rather than phenotypic measures of disease) as a proxy for psychiatric disease risk can be a useful alternative for research questions that would traditionally require long-term follow up and big datasets due to the low prevalence of some of the psychiatric disorders of interest in the population (e.g. adult-onset health consequences of child neurodevelopmental disorders). In addition, while high genetic risk for a psychiatric disorder is not always indicative of a diagnosis of psychiatric disease, it can index underlying subthreshold symptomatology that can still impact later adversities and quality of life (6).

So far, studies have used hypothesis-driven approaches to investigate associations of genetic risk for psychiatric disorders with various psychiatric and health outcomes as well as lifestyle factors (7, 8). However, big data resources that are readily available, such as UK Biobank with about 500,000 participants, provide rich phenotypic information that can be used for hypothesis-free studies and offset the multiple testing burden. Phenome scans are a type of hypothesis-free analysis where the association of a trait of interest is systematically tested with a potentially large number of phenotypes and can be hypothesis-generating by identifying an association when there is no prior reason to expect that an association may exist. As all available phenotypes are tested and the less ‘significant’ results published alongside those of greater ‘significance’, phenome scans can help to reduce biases associated with hypothesis-driven studies where researchers might only publish the most desirable or expected results.

In a Mendelian randomization phenome-wide association study (MR-pheWAS) genetic risk is used as a proxy for lifelong liability for a disorder to explore associations of this genetic liability with traits that may evolve as a consequence. Understanding these associations will be essential to inform prevention or early intervention strategies. However, conclusions about causality are limited due to the low predictive power and high pleiotropic effects of genetic risk scores for psychiatric conditions (8).

The aim of this study was to investigate the associations between genetic risk for five common psychiatric disorders – attention-deficit/ hyperactivity disorder (ADHD), autism spectrum disorder (ASD), schizophrenia (SCZ), major depression (MDD) and bipolar disorder (BP) - with a wide range of socio-demographic, lifestyle, physical and mental health outcomes in UK Biobank, using the systematic hypothesis-free MR-pheWAS approach.

## RESULTS

In total 334,976 participants of white British ancestry in UK Biobank were included in this study with an average age of 56 (standard deviation [SD]=8) years. A descriptive overview of selected UK Biobank study sample characteristics is given in Figure 1A. The UK Biobank participants are known to be more educated and healthier than the average UK population which is reflected in the high percentage of people with a university degree (47%) and low prevalence of current smoking (10%) in the sample, which is comparable to the full UK Biobank release (9). Furthermore, 34% of participants reported to have seen a general practitioner and 11% a psychiatrist for nerves, anxiety, tension or depression but there are few cases of schizophrenia (n=132), ADHD (n=71), ASD (n=143) or bipolar disorder (n=439). An overview of UK Biobank phenotype categories is given in Figure 1B.

**Figure 1.**
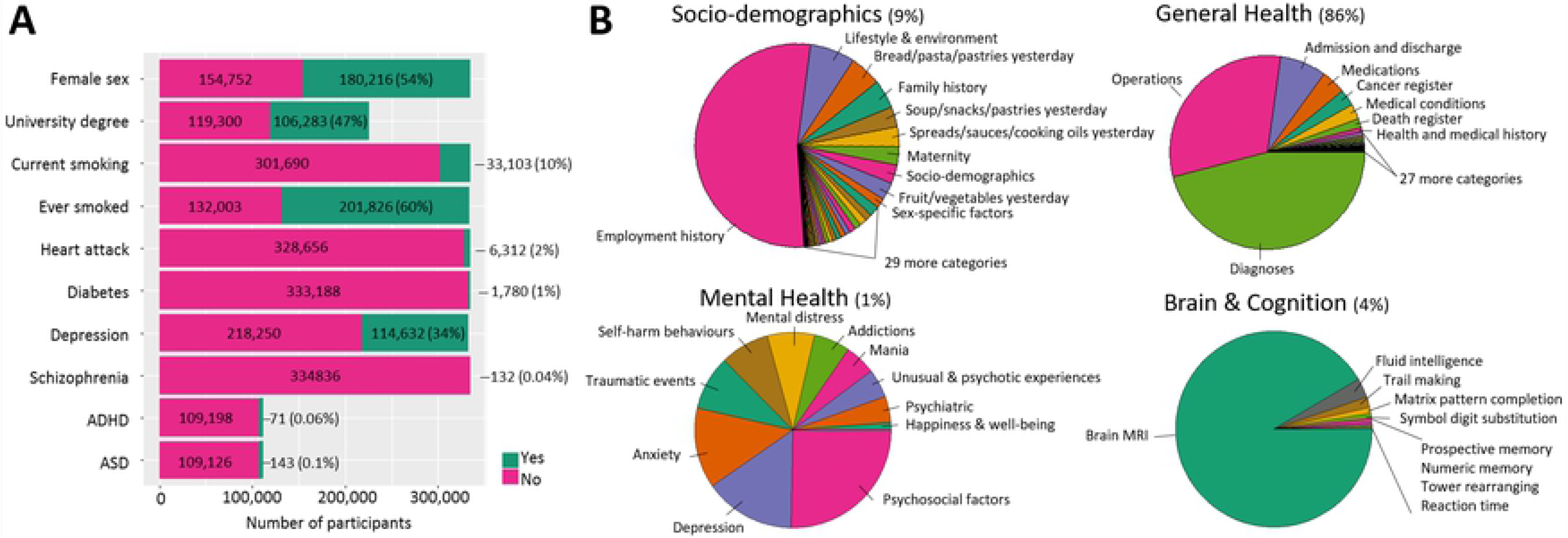
Study overview. (A) Descriptive sample overview of selected outcomes in UK Biobank. (B) Categories of UK Biobank with the size of pie chart sections indicating the number of included outcomes: socio-demographics (n=2,057), general health (n=19,740), mental health (n=233), brain and cognition (n=974).

## DISORDER SPECIFIC EFFECTS

The MR-pheWAS of each psychiatric disorder tested the association of the respective polygenic risk score, aggregated from independent, genome-wide significant SNPs, with 23,004 outcomes in UK Biobank, adjusted for age, sex and the first 10 genetic principal components. There was strong evidence after multiple testing correction based on the number of independent tests derived from spectral decomposition (p<2.55×10-6) for associations of either the ADHD, ASD, SCZ, MDD or BP PRS with 226 outcomes in 31 UK Biobank categories (Figure 2 and Table S1) as described below. A less stringent 5% FDR multiple testing threshold identified 209 additional outcomes also associated with at least one PRS (Figure S1). Correlations among the PRS can be found in supplementary Table S2. A detailed list of all MR-pheWAS results generated by the open-source PHESANT software package can be found in Table S3. Unless stated as a PHESANT result, estimates for continuous outcomes are generated by following up outcomes and manually curating the outcome phenotypes, to compute estimates on their original scale.

**Figure 2.**
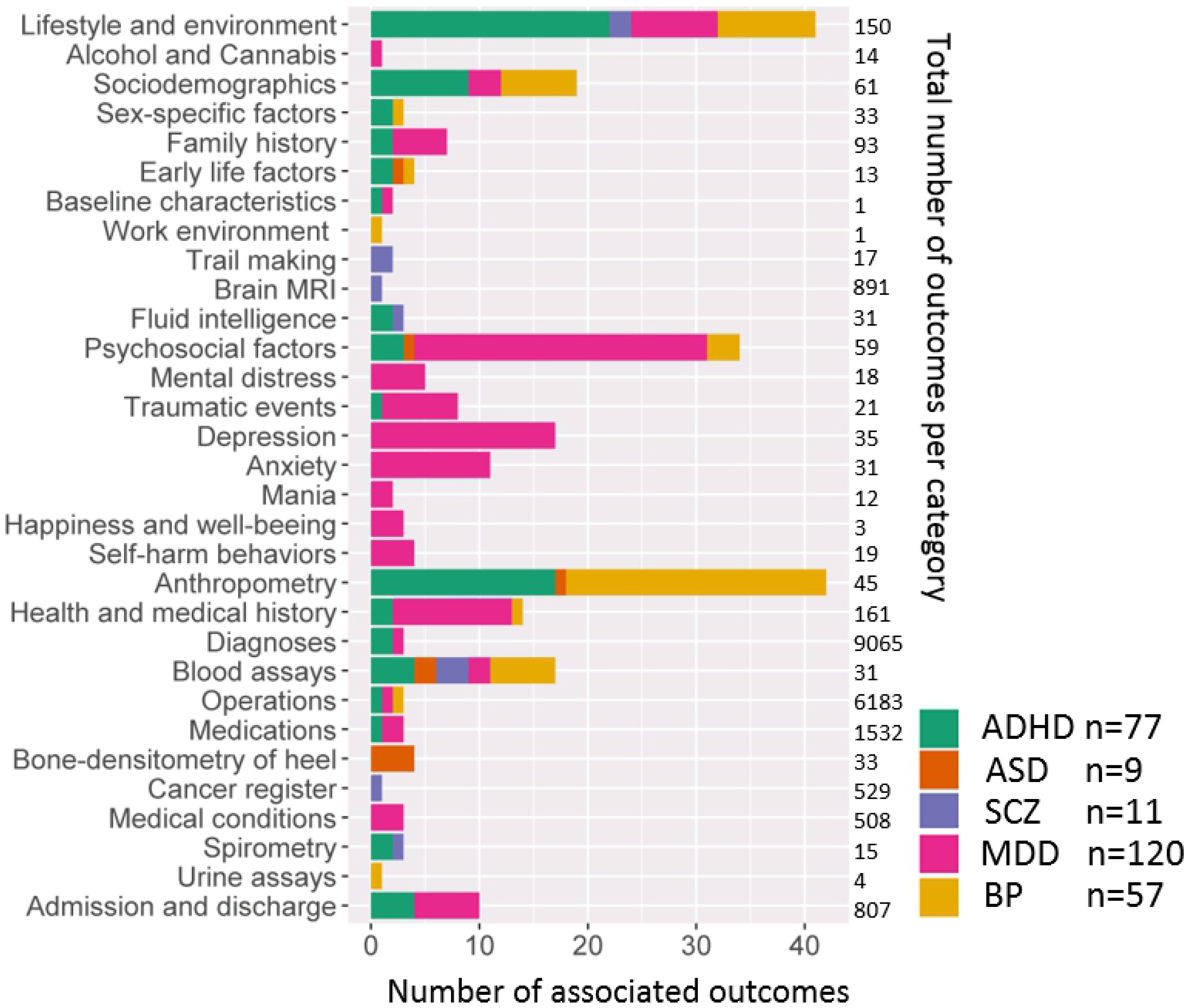
Overview of the distribution of disorder specific polygenic risk scores (p<5×10^−8^) associated outcomes per category of the UK Biobank variables catalogue. Shown are the number of associations with polygenic risk scores for attention deficit/hyperactivity disorder (ADHD), autism spectrum disorder (ASD), schizophrenia (SCZ), major depression (MDD) and bipolar disorder (BP).

### Attention deficit/ hyperactivity disorder

PRS_ADHD_ was strongly associated with 77 outcomes (Figure 3) including 39 socio-demographic factors, 33 general health and 5 mental health, brain and cognition outcomes. The strongest evidence of association with PRS_ADHD_ was seen for socio-demographic and lifestyle factors. A 1 SD higher PRS_ADHD_ was associated with a 1.09 month younger age at first sexual intercourse [95% confidence interval [CI]: −1.26,−0.94] (p=2.0×10^−16^), and 0.96 lower odds of having a university degree [95% CI: 0.95, 0.97] (p=1×10^−29^). In addition, PRS_ADHD_ was associated with younger maternal and paternal age of their parents (−0.08 years [95%CI: −0.103,−0.051] p=4.4×10^−9^; −0.10 years [95% CI: −0.134,−0.067] p=3.2×10^−9^,respectively), 0.97 lower odds of average household income [95%CI: 0.96,0.97] (p=1.3×10^−20^), 1.05 higher odds of current smoking [95%CI:1.03,1.06] (p=5.7×10^−15^) and 1.04 higher odds of experiencing physical abuse as a child [95%CI: 1.02,1.06] (p=1.3×10^−6^).

**Figure 3.**
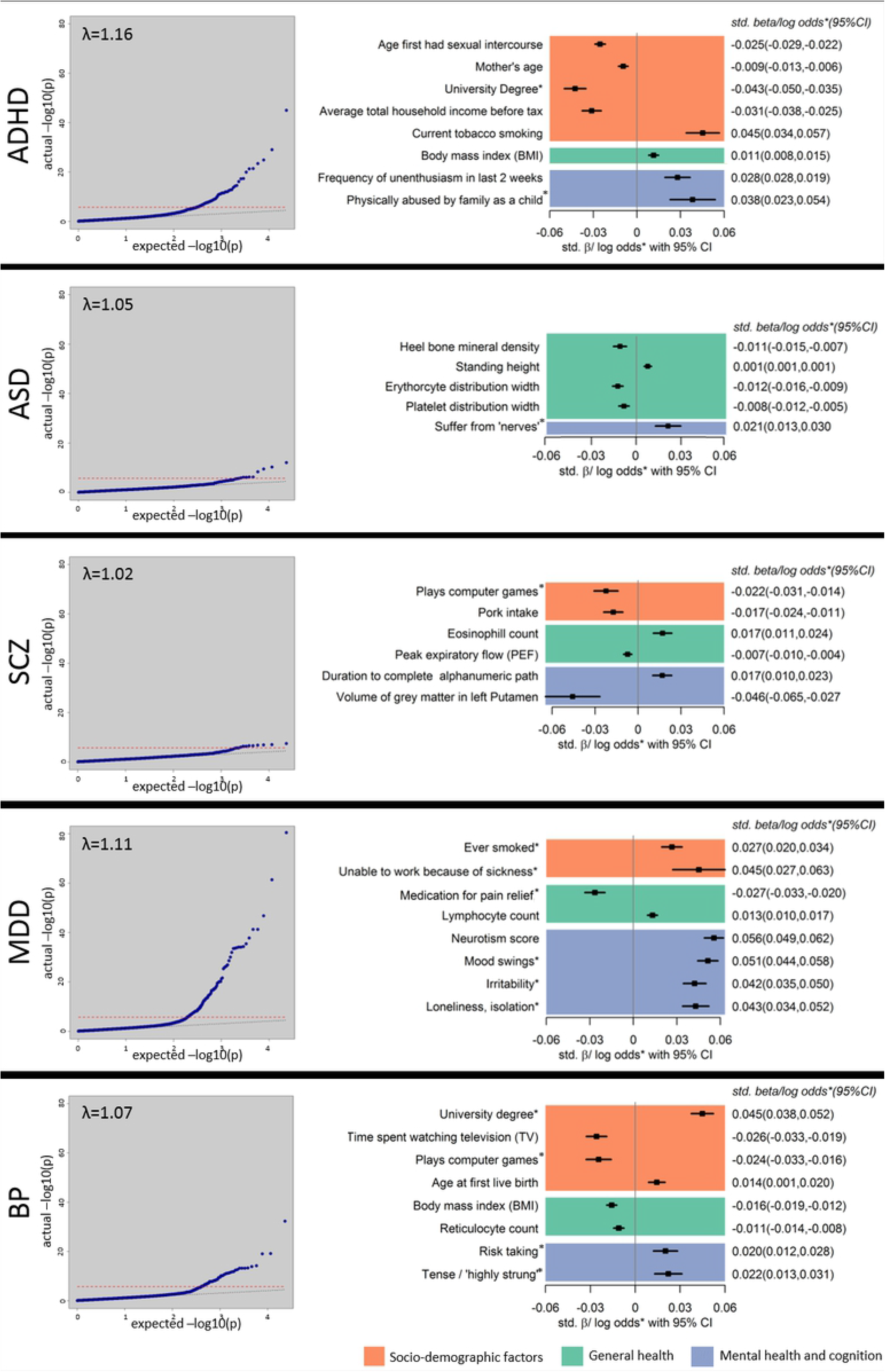
MR-PheWAS results for attention deficit/hyperactivity disorder (ADHD), autism spectrum disorder (ASD), schizophrenia (SCZ), major depressive disorder (MDD) and bipolar disorder (BP). Left hand panel: QQ plots of expected versus observed p-values for association of PRS with all outcomes in UK Biobank. Red line indicates the significance threshold (2.5×10^−6^). Lambda (λ) indicates the degree of inflation from the expected fit. Right hand panel: selected results from different categories with *p*-values below the significance threshold and estimates generated by PHESANT. Results for continuous outcomes (std. β) are the standard deviation change of inverse-rank normal transformed outcome per 1 SD higher PRS.

Further, 1 SD increase in PRS_ADHD_ was associated with 17 physical health outcomes related to obesity, including 0.05 kg/m^2^ higher BMI [95%CI: 0.037,0.070] (p=7.4×10^−11^), leg and arm fat mass, waist circumference and trunk fat mass. Furthermore, there was evidence for an association of PRS_ADHD_ with blood measures, such as 0.02 cells/L higher leukocyte count [95%CI: 0.012,0.026] (p=1.4×10^−7^).

Associations seen for brain and cognition include 0.04 points lower fluid intelligence score [95%CI: −0.050,−0.026] (p=1.9×10^−9^).

### Autism spectrum disorder

PRS_ASD_ was strongly associated with 9 outcomes (Figure 3), including 1 socio-demographic, 7 general health and 1 mental health outcome.

The strongest association of PRS_ASD_ was found for lower erythrocyte distribution width where 1 SD higher PRS_ASD_ was associated with 0.01% lower erythrocyte distribution width [95% CI: −0.013, −0.007] (p=6.3×10^−10^) and 0.98 lower odds of comparative body size at age 10 [95%CI:0.97,0.98] (p=6.2×10^−11^). Furthermore, 1 SD higher PRS_ASD_ was associated with 0.001 g/cm^2^ lower heel bone mineral density (BMD) [95%CI:−0.002,−0.001] (p=4.1×10^−5^).

The only mental health outcome that was associated with PRS_ASD_ was 1.02 higher odds of being a nervous person (“suffer from nerves”) [95%CI:1.01,1.03] (p=7.7×10^−7^).

### Schizophrenia

There was strong evidence of association for PRS_SCZ_ with 11 outcomes (Figure 3), including 2 socio-demographic, 4 mental health and cognition and 5 general health outcomes.

The strongest evidence of an association with PRS_SCZ_ was detected for playing computer games (OR:0.98 [95%CI:0.976,0.989] p=1.4×10^−7^), 0.01% lower platelet distribution width [95%CI:0.003,0.006] (p=1.6×10^−7^) and having 0.68 lower odds of glioma cancer [95%CI:0.592,0.785] (p=1.2×10^−7^).

In addition, 1 SD increased PRS_SCZ_ was associated with 0.42 sec longer duration of completing an online cognitive function test (alphanumeric path) [95%CI:0.237,0.580] (p=3.0×10^−6^) and a 19.1 mm^3^ reduced grey matter volume of the left putamen [95%CI:−26.9,−11.3] (p=1.5×10^−6^).

### Major depressive disorder

PRS_MDD_ was associated with 120 outcomes (Figure 3), including 18 socio-demographic, 76 mental health and 26 general health outcomes.

Most of the associations (63%) were related to mental health, including an association of higher PRS_MDD_ with higher odds of depression, anxiety, irritability, nervousness and mood swings. Strongest evidence of association with PRS_MDD_ was found for 1.07 higher odds of “seen a doctor for nerves, anxiety, tension or depression” [95%CI:1.066,1.081] (p=2.6×10^−81^), 0.11 points higher neuroticism score [95%CI:0.094,0.118] (p=2.0×10^−16^) and 1.05 higher odds of having mood swings [95%CI:1.045,1.060] (p=1.8×10^−47^).

Furthermore, there was strong evidence of 1 SD higher PRS_MDD_ being associated with socio-demographic and lifestyle traits including 1.03 higher odds of ever smoking [95%CI:1.02,1.03] (p=8.4×10^−14^) and 1.04 higher odds of cannabis use [95%CI:1.03,1.06] (p=2.3×10^−8^).

Associated physical health measures included 0.97 lower odds of taking medication for pain relief, constipation or heartburn [95%CI:0.967,0.981] (p=4×10^−6^) and 1.02 odds of more frequent feelings of pain, e.g. back pain [95%CI:1.015,1031] (p=6×10^−9^).

### Bipolar disorder

PRS_BP_ was associated with 57 outcomes (Figure 3), including 19 socio-demographic, 35 general health and 3 mental health outcomes.

Socio-demographic and lifestyle factors included associations of higher PRS_BP_ with 1.04 higher odds of having a university degree [95%CI:1.033,1.048] (p=4.5×10^−26^), 0.02 hours/day less time spent watching television [95%CI:−0.027,−0.017](p=2.8×10^−15^) and lower average alcohol intake (OR:0.98 [95%CI:0.978,0.989] p=1.5×10^−7^).

General health traits included 19 traits indicating an association of 1 SD higher PRS_BP_ with 0.07kg/m^2^ lower body weight and fat mass [95%CI:−0.090,−0.058] (p=2.0×10^−16^) and 5 traits related to blood measures, such as 0.004% decreased platelet distribution width [95%CI:−0.006,−0.003] (p=1.8×10^−6^).

The two traits related to mental health were risk taking (OR:1.03 [95%CI:1.018,1.035] p=2.9×10^−19^) and feeling fed-up (OR:0.98 [95%CI:0.975,0.989] p=2.5×10^−7^).

## CROSS DISORDER CONSIDERATIONS

The highest overlap of associated outcomes of the univariable MR-pheWAS scans was seen for ADHD and BP with 11 socio-economic and lifestyle and 12 general health outcomes associated with both disorders (Figure 4). However, the majority of the associations are directionally opposite for ADHD and BP. For example, higher PRS_ADHD_ showed evidence for associations with lower educational attainment and higher BMI, whereas higher PRS_BP_ was associated with higher educational attainment and lower BMI. Only a higher risk of smoking initiation (“ever smoked”) was directionally consistent for both PRS_ADHD_ and PRS_BP_.

**Figure 4.**
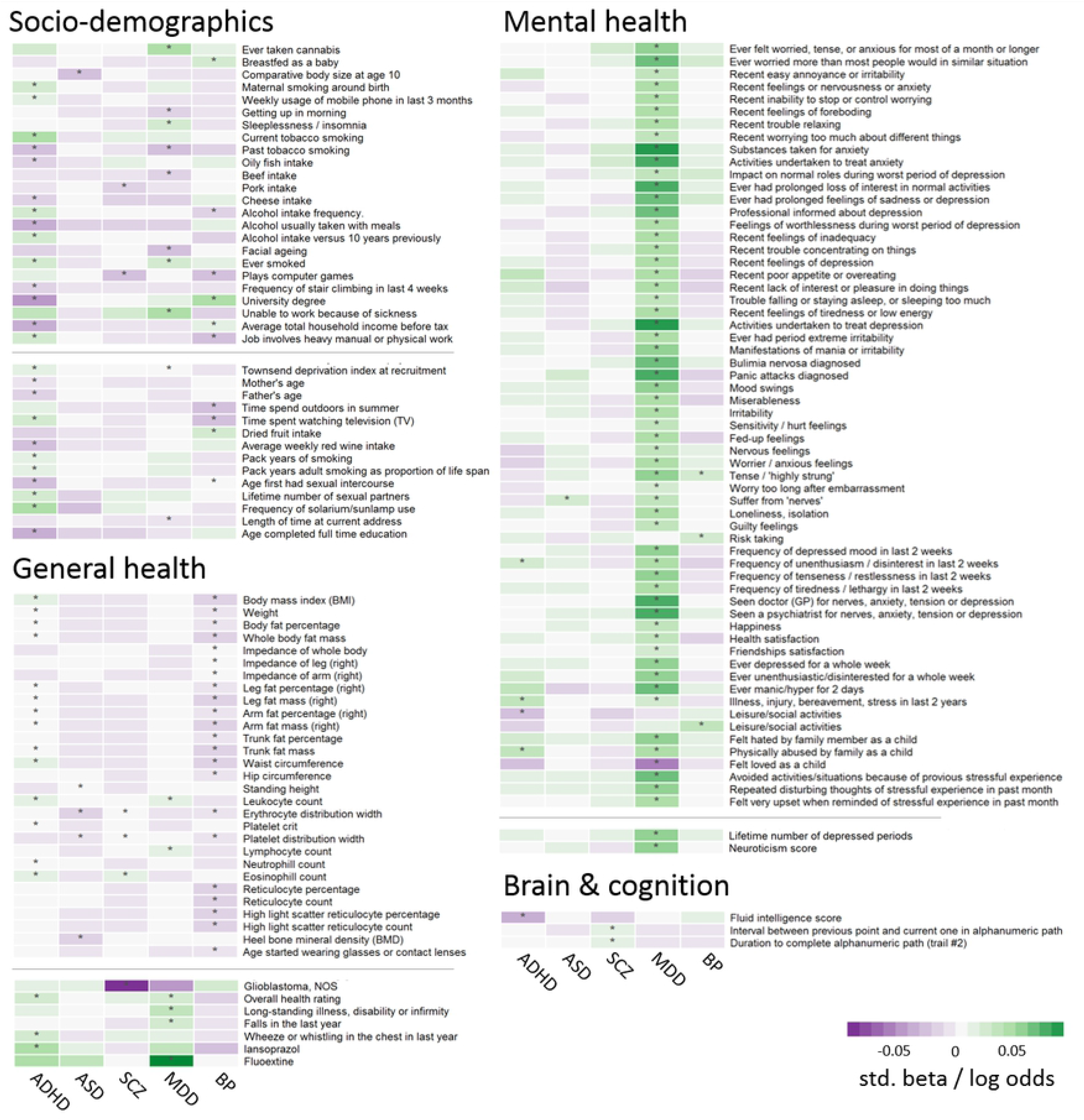
Cross-disorder comparison. Shown are standardized log odds (upper section in each panel) or standardized beta-values (lower section of each panel) of all outcomes associated with polygenic risk scores for either attention deficit/hyperactivity disorder (ADHD), autism spectrum disorder (ASD), schizophrenia (SCZ), major depressive disorder (MDD) or bipolar disorder (BP) at *p*<2.55×10^−6^ as indicated by stars (*). For outcomes categorized as ordered-logistic, only one outcome is displayed. Only associations with anthropometric measures of the right side of the body are shown. Estimates were generated by PHESANT. Results for continuous outcomes (std. beta) are the standard deviation change of inverse-rank normal transformed outcome per 1 SD higher PRS.

Furthermore, all disorder PRSs showed some evidence for association with different blood cell counts, such as a decreased leukocyte count for PRS_ADHD_ and PRS_MDD_, or a decreased eosinophil count for PRS_ADHD_ and PRS_SCZ_.

There was very little overlap of highly associated outcomes between the neurodevelopmental domains (ADHD, ASD and SCZ).

## SENSITIVITY ANALYSIS

We repeated our tests of association of outcomes passing the spectral decomposition threshold, additionally adjusting for additional potential confounders (assessment centre, genotype chip and the first 40 principal components). Estimates were highly consistent with our main results, as shown in Table S4.

Relaxing the p-value threshold for including SNPs in the PRS resulted in some changes in the results (Figure S2). For ADHD, SCZ, MDD and BP the general trend was an inflation of p-values, with more associations below the significance threshold (Table S5) and higher effect estimates with smaller confidence intervals. A different pattern was observed for autism spectrum disorder with inconsistent results for some of the outcomes, as described in detail in the supplementary Text S1. Overall the strength of associations obtained for blood cell count traits across disorders varied between p-value thresholds, with weaker associations found for less stringent p-value thresholds.

## DISCUSSION

In this study, we conducted a MR-pheWAS to examine the relationships between genetic liability for five major psychiatric disorders and 23,004 outcomes in about 300,000 UK Biobank participants.

Our results build on a large body of literature supporting links between genetic risk for psychiatric disorders with a wide variety of outcomes including psychological well-being, lifestyle, socio-demographic factors and physical health (2, 4, 7, 10, 11). Our findings also suggest that although different psychiatric disorders show strong genetic overlap (7), genetic risk for distinct psychiatric disorders show differential associations with lifestyle, socio-demographic factors and physical health as highlighted in Figure 5. Genetic liability for ADHD and bipolar disorder showed the strongest associations with lifestyle and social environmental factors as well as physical health. On the other hand, genetic liability for major depression was most strongly associated with psychological health and associations with lifestyle and socio-demographic factors were less robust.

**Figure 5.**
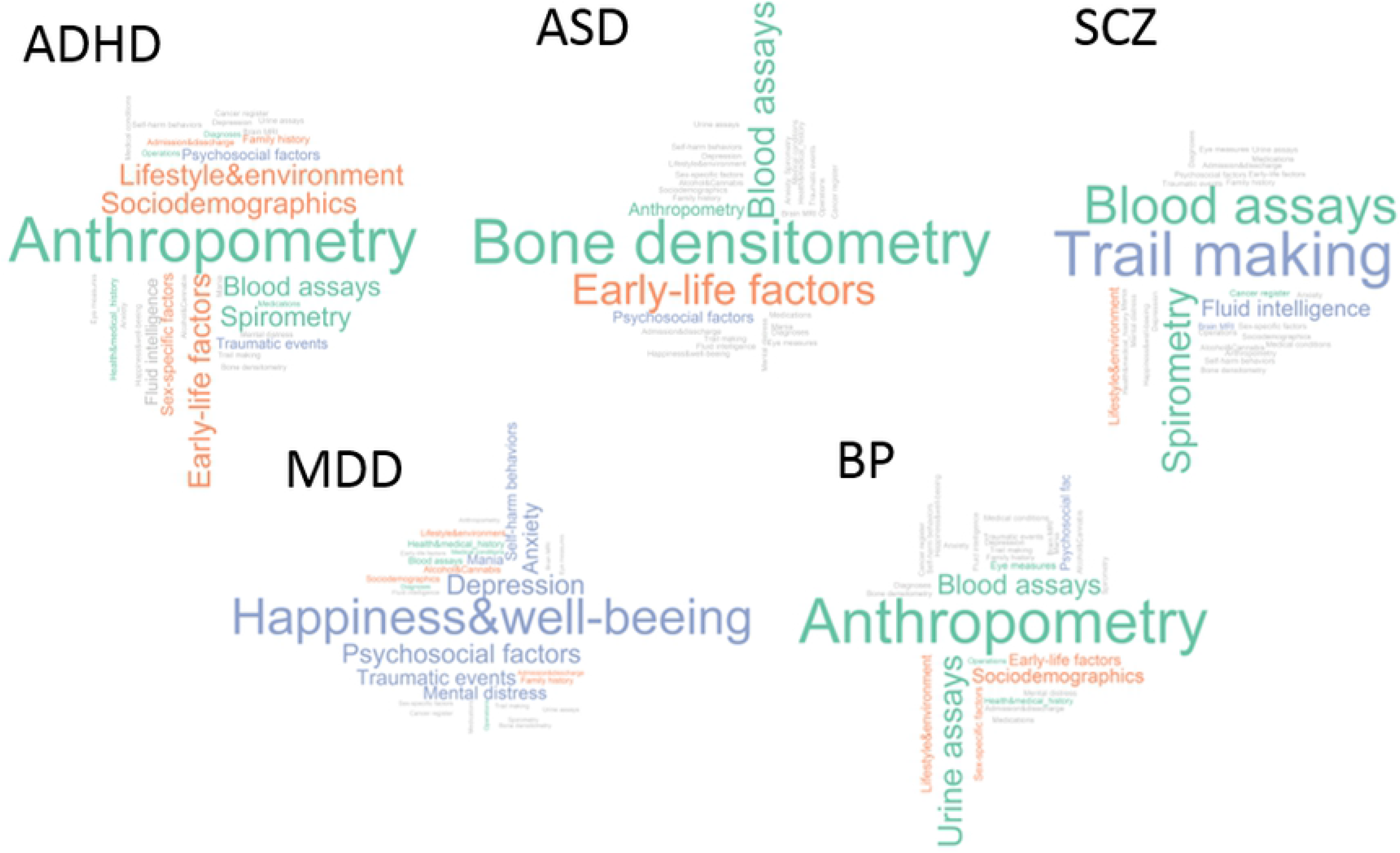
Categories of highly associated outcomes with polygenic risk scores for attention deficit/hyperactivity disorder (ADHD), autism spectrum disorder (ASD), schizophrenia (SCZ), major depressive disorder (MDD) and bipolar disorder (BP). Size of categories depends on the relative number of associated outcomes to the total number of outcomes within each category. Only categories with more than 1 variables are shown. Lifestyle and socio-demographic factors are shown in orange, physical health measures are shown in green and mental health, brain and cognition traits are shown in violet. Grey categories had zero hits for the corresponding disorder.

We were able to replicate previously reported associations between genetic liability for ADHD and lower educational attainment (12, 13), higher prevalence of smoking (14), younger age at delivery (15) and higher body mass index (16). While the previous findings for smoking and BMI were identified in young adults, our findings using an adult population-based sample with a mean age of 56 years, suggest that associations of childhood psychiatric disorder genetic liabilities with health and social outcomes persist into later adulthood. Associations of genetic liability for ADHD in childhood could represent effects of childhood ADHD or sub-threshold ADHD on long term social and economic outcomes, or alternatively associations could be due to parental effects (due to their shared genetic risk of ADHD) or horizontal pleiotropy (the same genetic variants affecting multiple traits).

Many of the associations of genetic liability for MDD with increased mood swings, irritability, feelings of loneliness and isolation are clinically known and have previously been reported (5). Our results are also in line with a recent publication from the Brainstorm consortium investigating genetic correlations among psychiatric disorders with neurological and quantitative traits using LD score regression and GWAS summary statistics, reporting high genetic correlations between most psychiatric disorders and educational attainment and BMI (2, 7). However, we found little evidence for associations of genetic liability for ADHD, ASD and schizophrenia with mental health outcomes, such as depressive symptoms, neuroticism or anxiety; and very few associations with cognitive or brain imaging outcomes, which might be because of the UK Biobank being a selected sample with lower rates of psychiatric disorders than the general population as discussed in the limitations section.

In addition to identifying previously reported associations of genetic liability for ASD, our MR-pheWAS also revealed novel associations. We found a strong association of genetic liability for ASD with decreased heel bone mineral density, which furthers previous evidence from observational studies that children and adolescents with ASD have lower bone mineral density (17, 18), higher frequency of bone fractures (19) and lower vitamin D levels (20, 21), which is essential for bone metabolism. This might suggest that these observed associations may be due to pleiotropic effects of genetic variants associated with bone health.

Our MR-pheWAS of schizophrenia suggested that genetic liability for schizophrenia was associated with lower grey matter volume in the left putamen and a lower risk for glioblastoma cancer. Both phenotypes have been associated with schizophrenia in observational studies but it is not clear whether these phenotypes are determinants or consequences of schizophrenia, or due to confounding or shared genetic risk (22, 23). For glioblastoma our finding could be attributed to common underlying mechanisms that act in opposite directions, since it has been previously suggested that the same biological pathways leading to schizophrenia may be protective for developing glioblastoma (23). With respect to differences in brain volumes of schizophrenia patients, two large studies by the international ENIGMA (24) and Japanese COCORO (25) consortia found no notable difference in putamen volume between schizophrenia patients and controls or associations of brain volumes with a PRS_SCZ_ in schizophrenia patients (26). Our results suggest that genetic risk for schizophrenia could be associated with putamen volume, but should be treated with caution because of potential selection bias due to the highly selected subset of about 10,000 UK biobank participants with brain scan data available at the time of data analysis (27). In line with our results, other previous work in schizophrenia patients and their relatives identified an association between schizophrenia and longer performance duration on the Trail Making Test (28), which requires searching and connecting irregularly arranged targets (digits and letters) in ascending order and is widely used to test for executive function, cognitive ability and processing speed (29–34).

Altered blood cell counts were associated with genetic liability for all disorders. Many psychiatric disorders previously have been associated with allergic or inflammatory states (35–37), such as asthma (38) and atopic diseases (39, 40) but it is unclear whether high inflammatory states are on the causal pathway to disorder manifestation or the result of comorbid and confounding behaviours associated with the disease, such as restricted diet, overweight, risky behaviours or medication. Our results support the possibility that altered blood cell counts could be a consequence of the disorder, but we cannot rule out contributions of horizontal pleiotropic effects. Also, considering the inconsistent findings from the sensitivity analyses for blood count traits, results need further validation and should be treated with caution.

### Limitations

Patients with psychiatric disorders or high genetic liability for psychiatric disorders are known to be less likely to participate in studies in the first place and more likely to drop-out during an ongoing study (41). Selection bias into a study as well as attrition can induce collider bias (42). There is consensus that the UK Biobank sample is not representative of the UK population, with participants showing, for example, lower prevalence of current smoking and lower rates of mortality (9). If both having a psychiatric disorder and a specific outcome (e.g. high socio-economic position) are associated with participation (the collider), this can induce an association between genetic risk for psychiatric disorders and the outcome, called collider bias. To reduce the possibility of collider bias we limited the set of included confounders in our main analysis but adjusted for assessment centre and genotype batch in sensitivity analysis.

A direct comparison of PHESANT estimates across the psychiatric disorders cannot be done without taking the differentially powered GWASs and derived PRS into account. This can also affect the number and set of outcomes associated with each disorder, which allows only a relative comparison among the outcomes. Further, the MDD GWAS used in the current study to calculate genetic risk scores included thirty thousand participants from UK Biobank (about 10% of the GWAS sample) which might have inflated our results for depression related items but is not expected to introduce bias in any other traits, such as blood counts.

Although genetic risk scores were derived using variants associated at genome-wide significance level, they can still have horizontal pleiotropic effects on different disorders and traits. Hence, our reported associations cannot on its own inform about causality but should be followed up with other causally informative methods. We therefore encourage triangulation of results using other study designs (43, 44), such as two-sample MR, negative control or twin studies.

### Conclusion

We were able to show that genetic liability for five common psychiatric disorders are associated with distinct aspects of adult life, including socio-demographic factors, mental and physical health. This is evident even in individuals from the general population who do not necessarily present with a psychiatric disorder diagnosis. However, there was surprisingly little overlap of findings for the different psychiatric disorder genetic risk scores despite the high genetic and symptomatic overlap of psychiatric disorders, such as schizophrenia and bipolar disorder. Furthermore, we want to emphasize the benefit of using genetic instruments for hypothesis-generating efforts in the field of psychiatry.

## METHODS

### Study population

Between 2006-2010 UK Biobank recruited 503,325 men and women in the UK at ages 40-69 years. The cohort contains a large dataset including physical measurements, blood/urine/saliva samples, health and lifestyle questionnaires as well as genotype (https://www.ukbiobank.ac.uk/).

For 463,010 participants genotyping was performed using the Affymetrix UK BiLEVE Axiom array or Affymetrix UK Biobank Axiom^®^ array. Participants with non-white British ancestry (n=54,757) and 73,277 who have a kinship coefficient denoting a third-degree relatedness were removed from an already quality checked dataset (excluding participants with withdrawn consent, sex mismatch or sex aneuploidy) (45), resulting in a dataset containing 334,976 participants (Figure 6).

**Figure 6.**
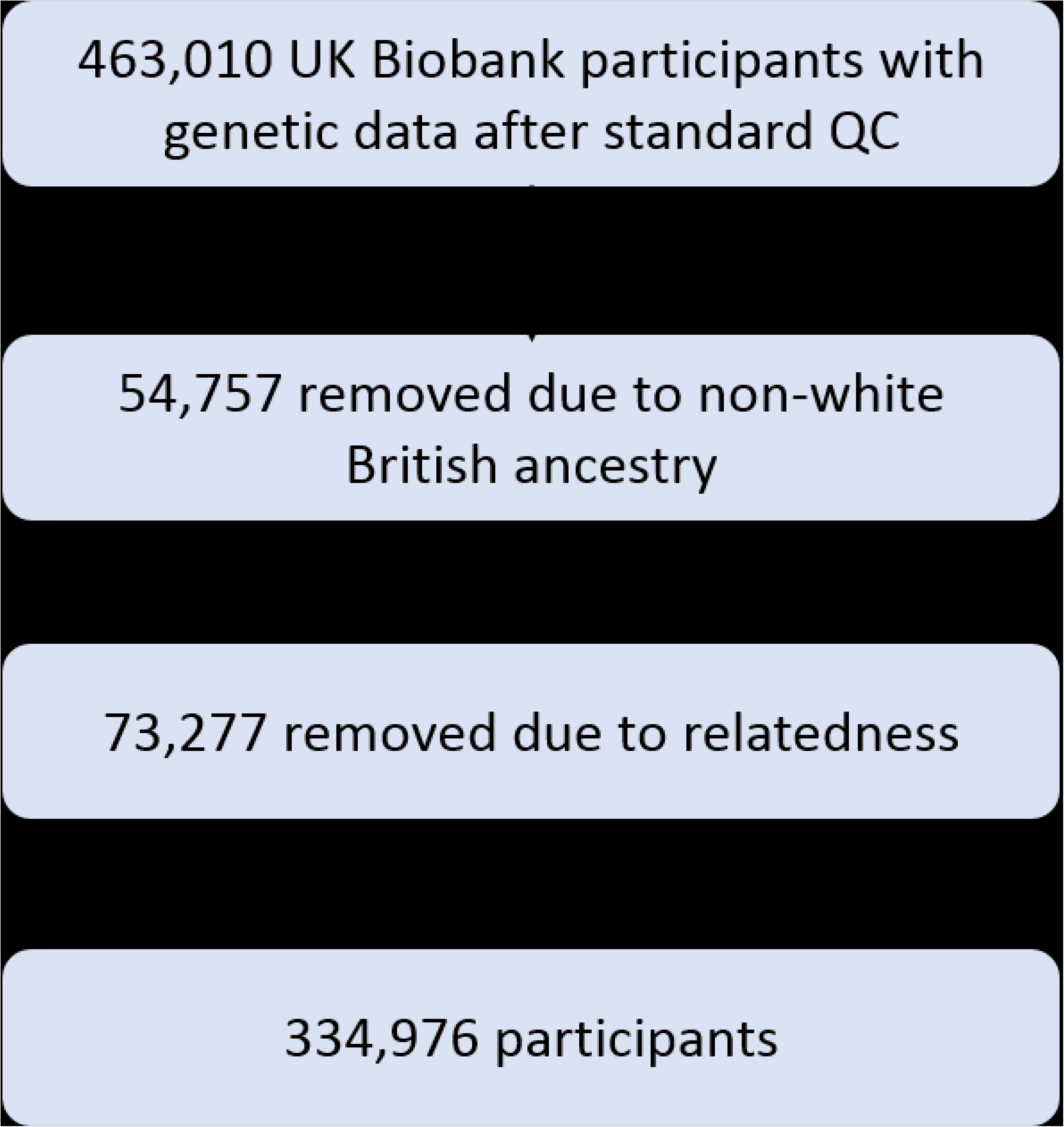
Overview of study sample derivation. Participants with withdrawn consent, sex mismatch or sex aneuploidy where already removed from the dataset in standard QC steps. (45)

UK Biobank received ethical approval from the research ethics committee (reference 13/NW/0382). All participants provided informed consent to participate. This work was done under application number 16729 (using genetic data version 2 [500K with HRC imputation] and phenotype dataset 21753).

### Polygenic risk scores

Genetic variants were identified from the most recent GWAS summary statistics listed in Table 1 with p<5×10^−8^ for ADHD, ASD, SCZ, MDD and BP. This stringent p-value cut-off was chosen to minimize bias introduced by horizontal pleiotropic effects of genetic variants. All summary statistics were subject to standard quality control including filtering for minor allele frequency (MAF>0.1) and imputation quality (INFO>0.8) and excluding the MHC region on chromosome 6 (26-33Mb) due to its complex linkage disequilibrium structure. Polygenic risk scores (PRS) were derived using PRSice v2.13 by identifying independent risk alleles in approximate linkage disequilibrium (R^2^<0.1 within 500kb distance) and computing a weighted, standardized mean score from these, as has been described previously (46).

**Table 1.** Details of GWAS used for calculating PRS

| *disorder* | *cases* | *controls* | *SNPs in PRS<sup>1</sup>* | *Source* |
| --- | --- | --- | --- | --- |
| ADHD | 20,183 | 35,191 | 10 | Demontis et al. (2019) (2) |
| ASD | 18,381 | 27,969 | 2 | Grove et al. (2017) (4) |
| Schizophrenia | 36,989 | 113,075 | 113 | Ripke et al. (2014) (3) |
| MDD | 135,458 | 344,901 | 34 | Wray et al. (2018) (5) |
| Bipolar disorder | 20,129 | 21,524 | 8 | Ruderfer et al. (2018)(47) |

1- PRS derived from genome-wide significant hits ( $p < 5 \times 10^{-8}$ )
2- SNP heritability estimates reported in the corresponding discovery sample
ADHD – Attention deficit/hyperactivity disorder, MDD – Major depression, ASD – Autism spectrum disorder

### Outcomes

UK Biobank provides a fully searchable data showcase (http://biobank.ctsu.ox.ac.uk/crystal/) which at the time of data download (March 2018) included 23,004 outcomes (see supplementary Text S2), including lifestyle and environment, socio-demographic, early life factors, anthropometry, family history and depression outcomes.

Age, sex and the first 10 principal components derived from the genetic data were included as covariates in all regression models. Age was derived from the participants date of birth and the date of their first assessment centre visit. Sex was self-reported and validated using genetic data.

### PHESANT MR-pheWAS

PHESANT package (version 0.17) was used to test the association of each PRS with each outcome variable in Biobank. A detailed description of PHESANT’s automated rule-based method is given elsewhere (48, 49). In brief, decision rules are based on the variable field type and categorize each variable as one of four data types: continuous, ordered categorical, unordered categorical or binary. PHESANT then estimates the univariate association of the PRS (independent) with each outcome variable (dependent) in a regression model, respectively. Normality of continuous data is ensured by an inverse normal rank transformation prior to testing. All estimates correspond to 1 SD change of the PRS. Selected continuous outcomes were followed up to compute meaningful estimates on the original phenotype scale for better interpretation.

PHESANT assigns each UK Biobank outcome to one of 91 level 3 categories based on the 235 origin categories of the UK Biobank catalogue (a full list of categories is provided in Table S1). Furthermore, three authors (BL, EW, ES) grouped these 91 categories into four prespecified higher level categories in order to aid result presentation: socio-demographics and lifestyle, brain and cognition, mental health and general health (Figure 1B).

To account for multiple testing (n=23,004 tests) we used a previously derived threshold (49, 50) based on an estimate of the number of independent phenotypes calculated using spectral decomposition (phenoSPD) (n=19,645). The multiple testing adjusted significance threshold was p<2.55×10^−6^ (0.05/19,645). The amount of inflation of observed versus expected p-values is given as the ratio of the median chi-squared statistics for observed to expected median p-values, referred to as Lambda (λ). A conservative Bonferroni correction of multiple testing that assumes uncorrelated traits, would yield a similar p-value threshold of p<2.30×10^−6^ (0.05/23,004).

### PHESANT sensitivity analysis

Analyses were re-run to assess residual confounding of assessment centre and genetic batch, including them as well as all 40 principal components as additional covariates for outcomes identified as strongly associated with either one of the disorders PRS. These covariates were not included in the first model because this could introduce collider bias if, for example, location of assessment centre is affected by both genetic predisposition and outcomes, as discussed in the limitations section.

Furthermore, PRS were derived using various p-value thresholds (p<0.01, p <0.1×10^−3^, p<1×10^−4^, p<1×10^−5^, p<1×10^−6^) with consequently increasing numbers of SNPs (Table S6) and the five MR-pheWAS were re-run with the more relaxed PRS to capture a larger amount of explained variation in the disorders by accepting an increase in horizontal pleiotropic effects. For MDD GWAS results were available for only 10,000 SNPs at these additional thresholds due to availability restrictions.

All analyses were performed in R version 3.2.4 ATLAS and R version 3.3.1, and the code is available at [https://github.com/MRCIEU/Psychiatric-disorder-pheWAS-UKBB]. Git tag v0.1 corresponds to the version presented here.

## ACKNOWLEDGEMENTS

This research was conducted using the UK Biobank resource under application number 16729. We acknowledge the members of the Psychiatric Genomics Consortium (PGC) and The Lundbeck Foundation Initiative for Integrative Psychiatric Research (iPSYCH) for the publicly available data used as the discovery samples in this article. Further we want to acknowledge Richard Anney for providing us the quality controlled GWAS summary statistics and Mick O’Donovan for his advice regarding the use of GWAS summary statistics.

## AUTHOR CONTRIBUTIONS

BL: Data curation, Formal analysis, Investigation, Methodology, Software, Visualization, Writing – original draft, Writing – review & editing

LACM: Methodology, Software, Writing – review & editing

LR: Writing – review & editing

GDS: Methodology, Writing – review & editing

AT: Conceptualization, Investigation, Funding acquisition, Writing–review & editing

KT: Conceptualization, Methodology, Writing–review & editing

EW: Conceptualization, Investigation, Methodology, Writing–review & editing

ES: Conceptualization, Investigation, Methodology, Writing–review & editing

## FUNDING

BL and LR are supported by the Wellcome Trust (grant ref: 204895/Z/16/Z) awarded to AT, GDS, ES and KT. BL, LACM, GDS, KT, EW and ES work in a unit that receives funding from the University of Bristol and the UK Medical Research Council (MC_UU_00011/1 and MC_UU_00011/3). LACM is funded by a University of Bristol Vice-Chancellor’s Fellowship.

## CONFLICT OF INTEREST

The authors declare no conflict of interest.

## SUPPORTING INFORMATION

**Figure S1**. Overview of the distribution of disorder specific polygenic risk scores (p<5×10^−8^) associated outcomes per category of the UK Biobank variables catalogue after FDR adjustment for multiple testing. Shown are the number of associations with polygenic risk scores for attention deficit/hyperactivity disorder (ADHD), autism spectrum disorder (ASD), bipolar disorder (BP), major depressive disorder (MDD) and schizophrenia (SCZ).

**Figure S2**. Top associated outcomes with PRS for attention deficit/hyperactivity disorder (ADHD), autism spectrum disorder (ASD), schizophrenia (SCZ), bipolar disorder (BP) and major depressive disorder (MDD) across different p-value thresholds for SNP inclusion (5×10-8 − 1×10-2).

**Table S1**. Overview of UK Biobank categories with total number of outcomes per category and number of associated outcomes with polygenic risk scores passing the significance threshold (p<2.55×10^−6^). (ADHD-attention defict/ hyperactivity disorder, ASD-autism spectrum disorder, SCZ-schizophrenia, MDD-major depressive disorder, BP-bipolar disorder)

**Table S2**. Correlation matrix of polygenic risk scores (p<5×10^−8^). Correlation coefficients are displayed on the left side, p-values on the right side of the table. (ADHD-attention defict/ hyperactivity disorder, ASD-autism spectrum disorder, SCZ-schizophrenia, MDD-major depressive disorder, BP-bipolar disorder)

**Table S3**. MR-PheWAS results for association of generic risk of 5 common psychiatric disorders with 23,004 outcomes in UK Biobank. Genetic risk is defined as weighted sum of all genome-wide significant risk alleles for each disorder in 334,976 participants of the UK Biobank. Estimates were generated by PHESANT. Results for continuous outcomes are the standard deviation change of inverse-rank normal transformed outcome per 1 SD higher PRS.

**Table S4**. MR-PheWAS follow-up and sensitivity results for selected continuous outcomes. Genetic risk is defined as weighted sum of all genome-wide significant risk alleles for each disorder in 334,976 participants of the UK Biobank. Estimates were generated by linear regression on the original variable scale per 1 SD higher PRS.

**Table S5**. Number of strongly associated traits with PRS for attention-deficit/hyperactivity disorder (ADHD), autism spectrum disorder (ASD), schizophrenina (SCZ), major depressive disorder (MDD) and bipolar disorder (BP) at different p-value thresholds for PRS calculation.

**Table S6**. Number of SNPs included in polygenic risk scores for attention-deficit/hyperactivity disorder (ADHD), autism spectrum disorder (ASD), schizophrenina (SCZ), major depressive disorder (MDD) and bipolar disorder (BP) at different p-value thresholds.

**Text S1**. Sensitivity analysis for autism spectrum disorder

**Text S2**. UK Biobank outcomes description

